## SupplementaryInformation for "Integrative genomics identifies a convergent molecular subtype that links epigenomic with transcriptomic differences in autism"

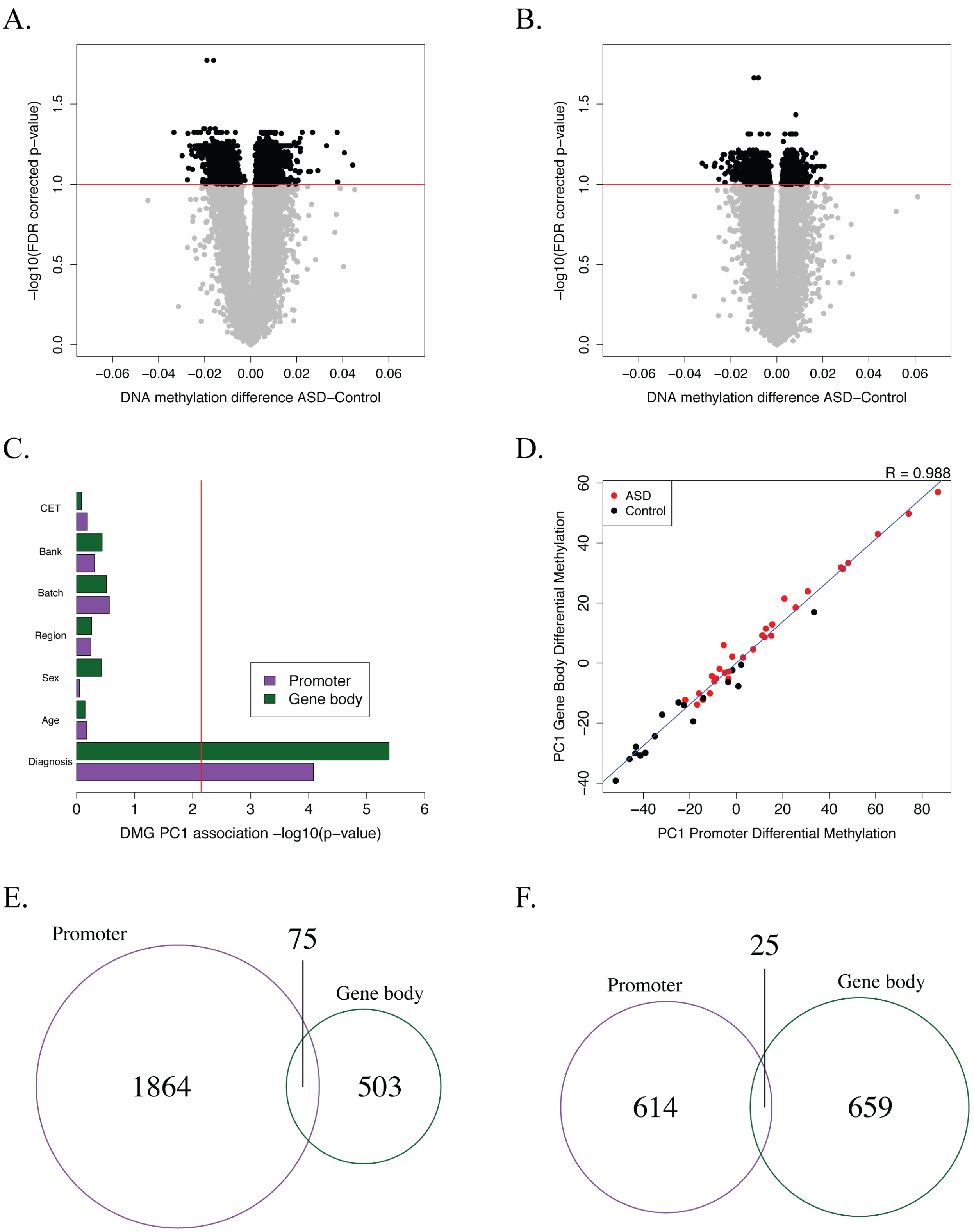

A.
B.
−log10(FDR corrected p−value)
DNA methylation difference ASD−Control
C.
CET
Bank
Batch
Region
Sex
Age
Diagnosis
Promoter
Gene body
DMG PC1 association −log10(p−value)
D.
R = 0.988
ASD
Control
PC1 Gene Body Differential Methylation
PC1 Promoter Differential Methylation
E.
Promoter
75
Gene body
1864
503
F.
25
Promoter
Gene body
614
659

**Supplementary Figure 1**. Initial differential DNA methylation analysis.

**A**) Volcano plot showing relationship between methylation difference and significance for differential promoter methylation analysis between all ASD samples and controls. A significance FDR cutoff of 0.1 is shown.

**B**) Volcano plot showing relationship between methylation difference and significance for differential gene body methylation analysis between all ASD samples and controls. A significance FDR cutoff of 0.1 is shown.

**C**) Relationship between sample loadings on the first principal component of differentially methylated promoters and gene bodies with technical and biological covariates. A Bonferroni corrected P-value of 0.05 is marked with a red line.

**D**) Relationship between sample loadings on the first principal component of differentially methylated promoters and gene bodies.

**E**) Overlap in genes between differentially methylated promoters and gene bodies that are hypermethylated in ASD.

**F**) Overlap in genes between differentially methylated promoters and gene bodies that are hypomethylated in ASD.

A.

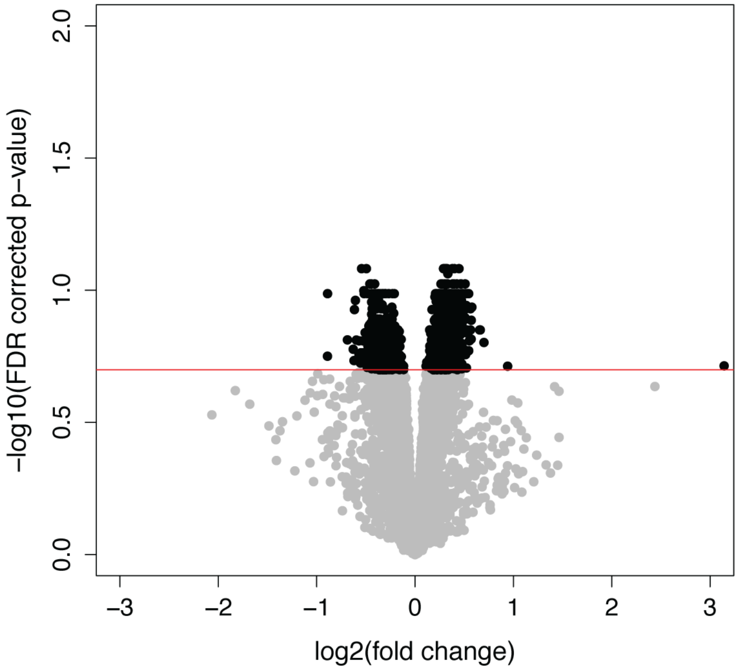
-log10(FDR corrected p-value)
2.0
1.5
1.0
0.5
0.0
-3
-2
-1
0
1
2
3
log2(fold change)

B.

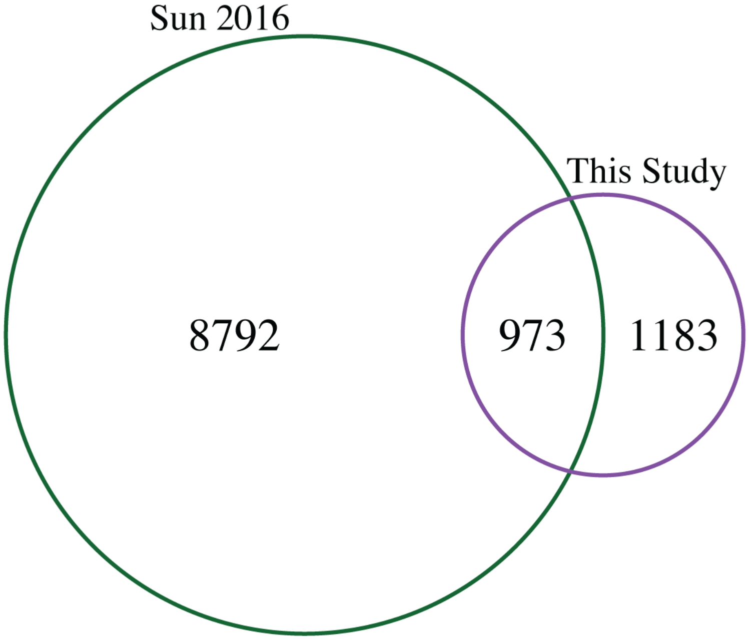
Sun 2016
This Study
8792
973
1183

**Supplementary Figure 2**. Initial differential histone acetylation analysis.

**A**) Volcano plot showing relationship between acetylation fold change and significance for differential histone acetylation analysis between all ASD samples and controls. A significance FDR cutoff of 0.2 is shown.

**B**) Overlap in differentially acetylated peaks between this analysis and Sun et al 2016.

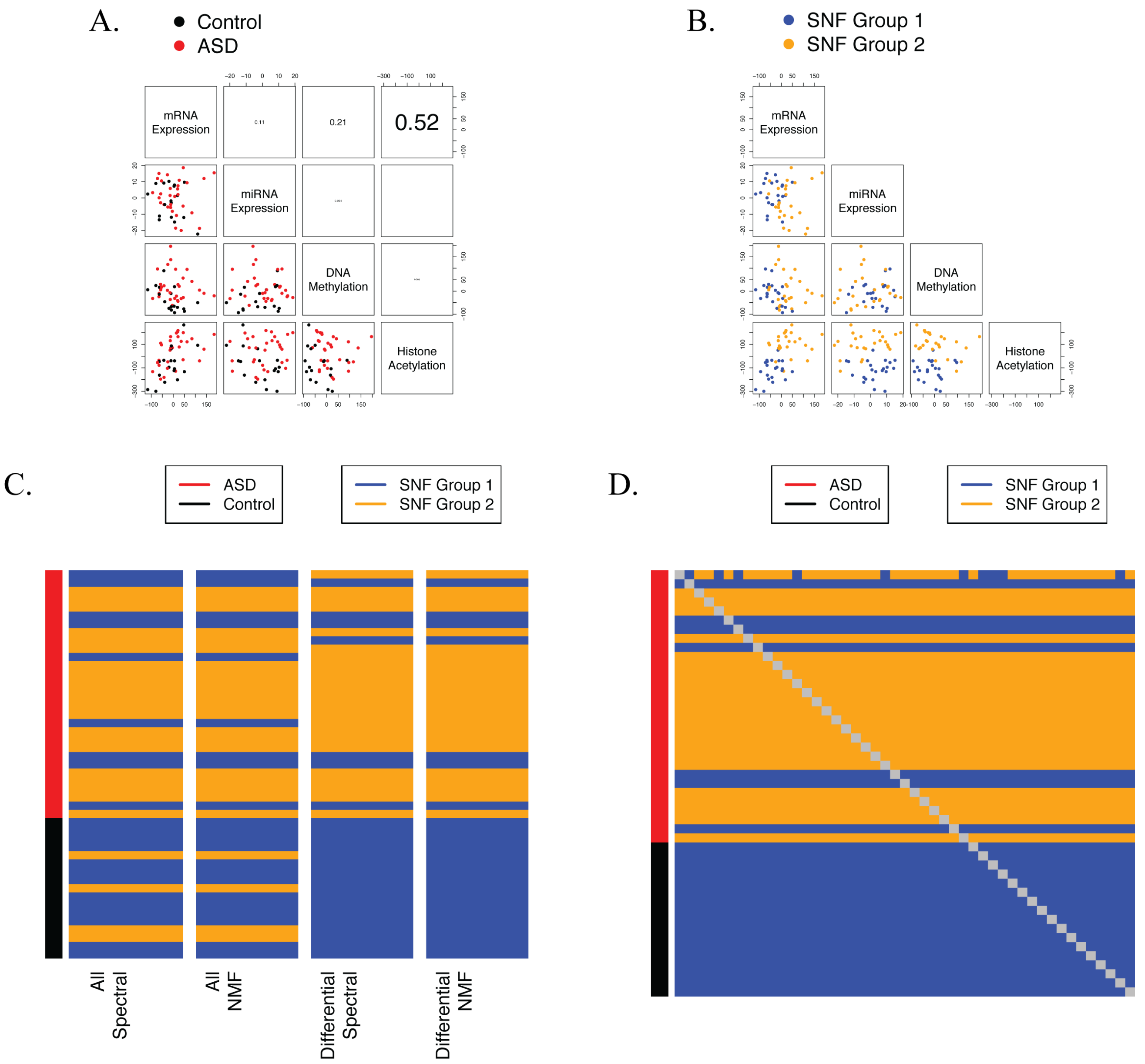
A.
Control
ASD
mRNA Expression
0.11
0.21
0.52
miRNA Expression
DNA Methylation
Histone Acetylation
B.
SNF Group 1
SNF Group 2
mRNA Expression
miRNA Expression
DNA Methylation
Histone Acetylation
C.
ASD
Control
SNF Group 1
SNF Group 2
All Spectral
All NMF
Differential Spectral
Differential NMF
D.
ASD
Control
SNF Group 1
SNF Group 2

**Supplementary Figure 3**. SNF sample clustering.

**A**) Relationship between sample loadings on the first principal component of mRNA expression, miRNA expression, DNA methylation, and histone acetylation when utilizing all features from each dataset.

**B**) Classification of 2 sample clusters using SNF on all features from each of the 4 datasets.

**C**) Comparison of SNF cluster assignments when using all features or restricting to differential features as well as when using spectral clustering or nonnegative matrix factorization (NMF).

**D**) Comparison of SNF cluster assignments when exhaustively leaving each sample out and performing clustering on the remaining samples.

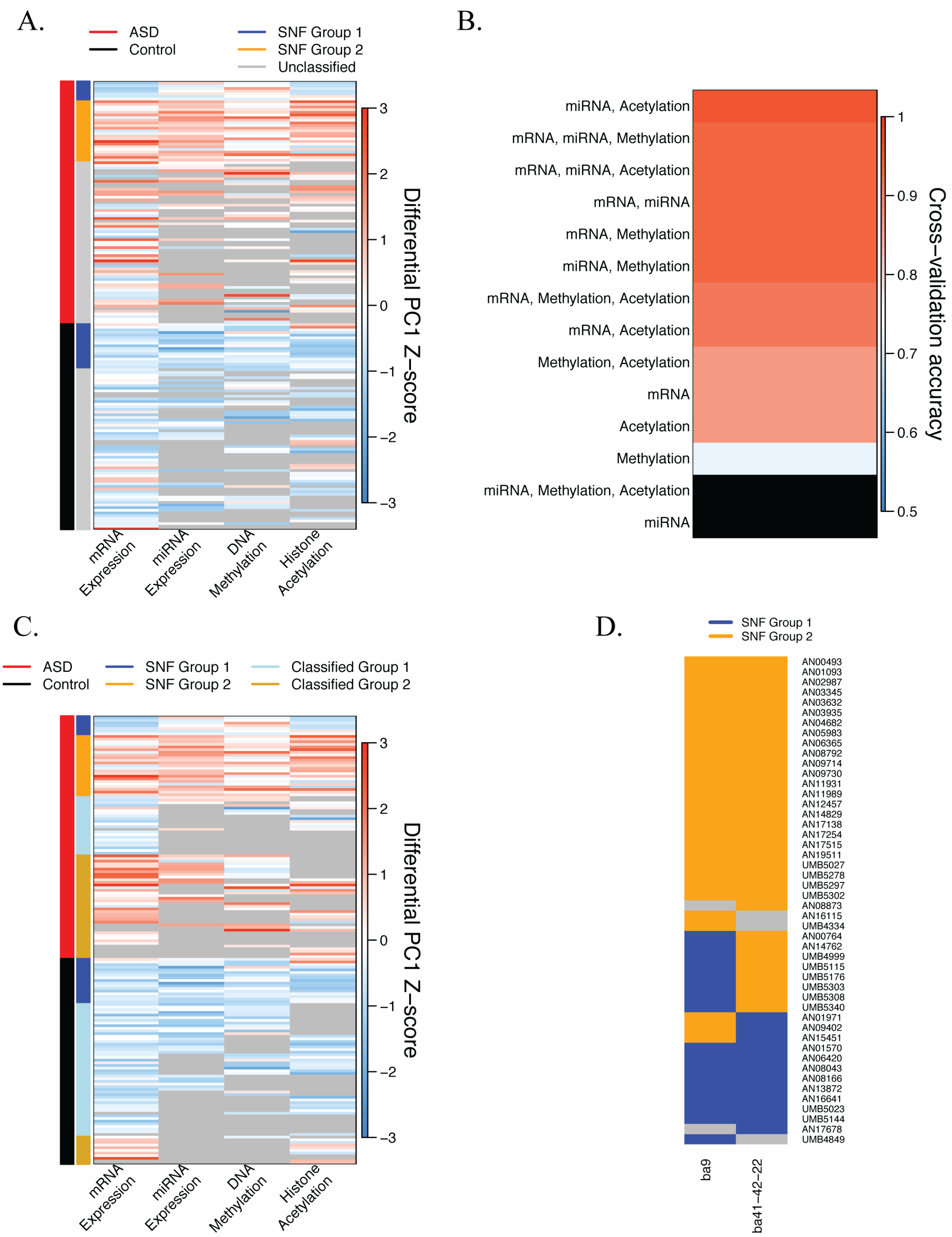
A.
ASD
Control
SNF Group 1
SNF Group 2
Unclassified
mRNA Expression
miRNA Expression
DNA Methylation
Histone Acetylation
Differential PC1 Z–score
3
2
1
0
−1
−2
−3
B.
miRNA, Acetylation
mRNA, miRNA, Methylation
mRNA, miRNA, Acetylation
mRNA, miRNA
mRNA, Methylation
miRNA, Methylation
mRNA, Methylation, Acetylation
mRNA, Acetylation
Methylation, Acetylation
mRNA
Acetylation
Methylation
miRNA, Methylation, Acetylation
miRNA
Cross–validation accuracy
1
0.9
0.8
0.7
0.6
0.5
C.
ASD
Control
SNF Group 1
SNF Group 2
Classified Group 1
Classified Group 2
mRNA Expression
miRNA Expression
DNA Methylation
Histone Acetylation
Differential PC1 Z–score
3
2
1
0
−1
−2
−3
D.
SNF Group 1
SNF Group 2
AN00493
AN01093
AN02987
AN03345
AN03632
AN03935
AN04682
AN05983
AN06365
AN08792
AN09714
AN09730
AN11931
AN11989
AN12457
AN14829
AN17138
AN17254
AN17515
AN19511
UMB5027
UMB5278
UMB5297
UMB5302
AN08873
AN16115
UMB4334
AN00764
AN14762
UMB4999
UMB5115
UMB5176
UMB5303
UMB5308
UMB5340
AN01971
AN09402
AN15451
AN01570
AN06420
AN08043
AN08166
AN13872
AN16641
UMB5023
UMB5144
AN17678
UMB4849
ba9
ba41–42–22

**Supplementary Figure 4**. Classification of samples into the 2 molecular subtypes.

**A**) Sample Z-score loadings for differential mRNA expression, miRNA expression, DNA methylation, and histone acetylation after running SNF, but before classification of samples that were missing in one or more datasets. The samples were sorted first by diagnosis, then by SNF subtype assignment. Samples not present in a particular dataset are colored in grey.

**B**) Exhaustive leave one out cross-validation accuracies for each logistic regression classification model. Two models (miRNA/Methylation/Acetylation and miRNA only) did not have any test samples and are colored black.

**C**) Sample Z-score loadings for differential mRNA expression, miRNA expression, DNA methylation, and histone acetylation after classification of samples that were missing in one or more datasets. The samples were sorted first by diagnosis, then by SNF subtype assignment. Samples not present in a particular dataset are colored in grey.

**D**) Sample subtype assignments for 2 cortical regions from 48 ASD individuals. Individuals not present in any of the 4 datasets from a particular cortical region are colored in grey.

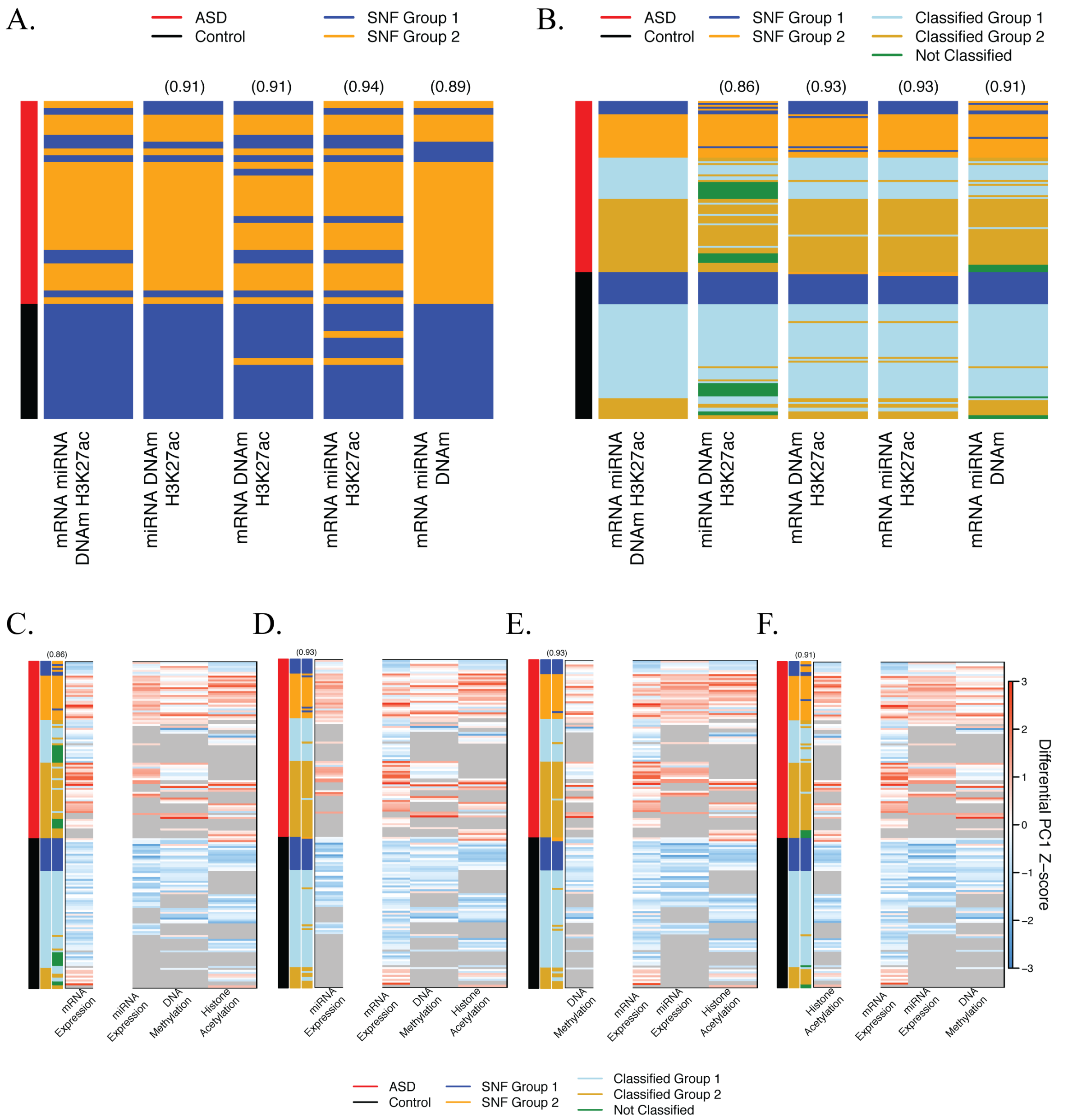
A.
ASD
Control
SNF Group 1
SNF Group 2
(0.91)
(0.91)
(0.94)
(0.89)
mRNA miRNA DNAm H3K27ac
miRNA DNAm H3K27ac
mRNA DNAm H3K27ac
mRNA miRNA H3K27ac
mRNA miRNA DNAm
B.
Classified Group 1
Classified Group 2
Not Classified
(0.86)
(0.93)
(0.93)
(0.91)
C.
D.
E.
F.
mRNA Expression
miRNA Expression
DNA Methylation
Histone Acetylation
Differential PC1 Z−score

**Supplementary Figure 5**. Robustness of sample classifications.

**A**) Comparison of SNF clustering assignments for 47 samples using all of the four datasets to those when utilizing three out of four datasets. The samples are sorted by diagnosis. The concordances of sample assignments to those when using the complete dataset are shown in parentheses. The datasets used in SNF clustering are shown below each column.

**B**) Comparison of SNF clustering and classification assignments for 169 samples using all of the four datasets to those when utilizing three out of four datasets. The samples are sorted by diagnosis. The concordances of sample assignments to those when using the complete dataset are shown in parentheses. The datasets used in SNF clustering are shown below each column.

**C**) Sample Z-score loadings for differential mRNA expression, miRNA expression, DNA methylation, and histone acetylation after SNF clustering and classification of samples using miRNA expression, DNA methylation, and histone acetylation datasets while holding out the mRNA expression dataset (third y-axis color label). The samples were sorted first by diagnosis (first y-axis color label), then by subtype assignment when using all four datasets (second y-axis color label). The concordance between subtype assignments when using all four datasets as compared to those when using only three datasets is shown in parenthesis.

**D**) Same as **C**, except the held-out dataset is miRNA expression.

**E**) Same as **C**, except the held-out dataset is DNA methylation.

**F**) Same as **C**, except the held-out dataset is histone acetylation.

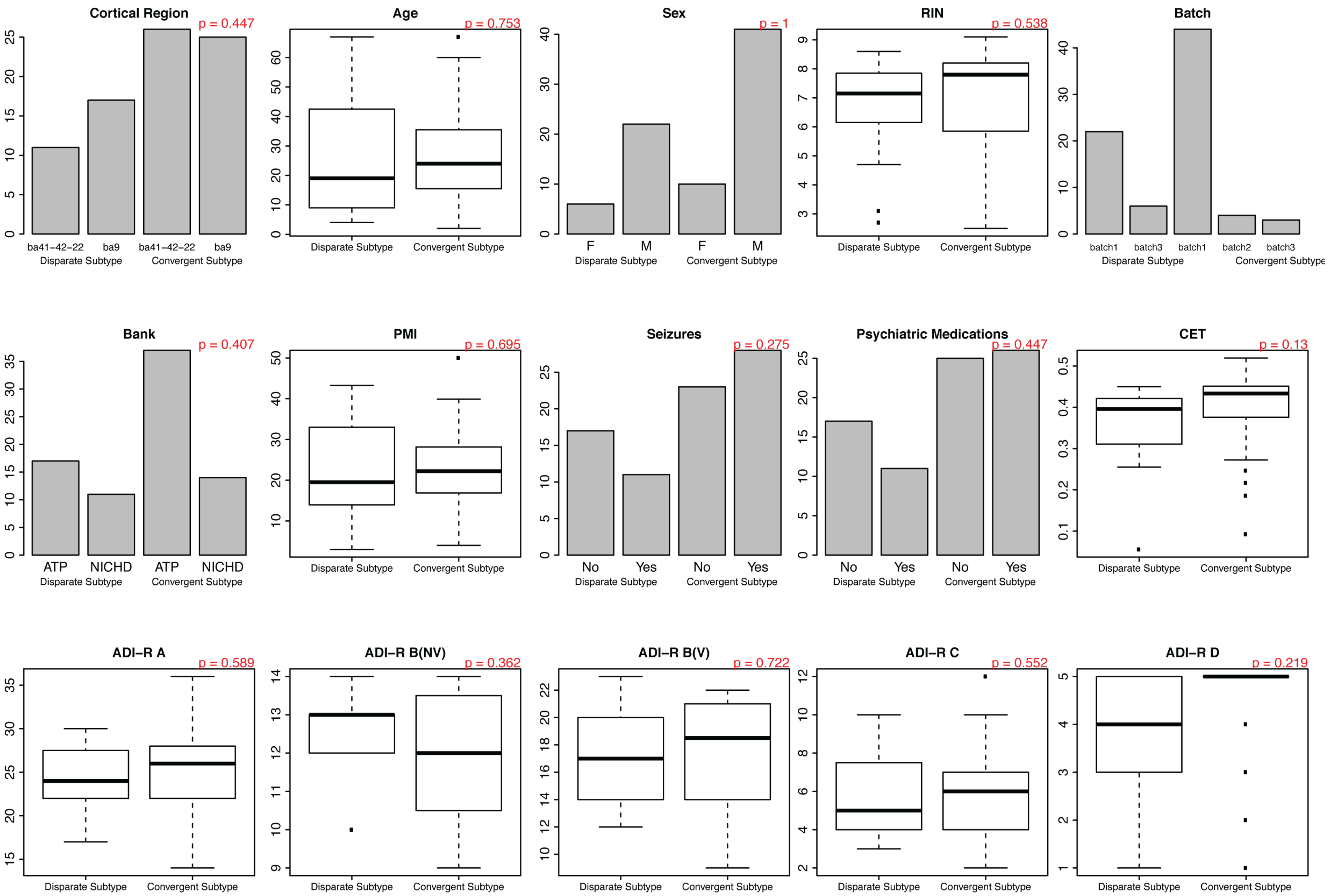

Cortical Region
p = 0.447
ba41−42−22
ba9
ba41−42−22
ba9
Disparate Subtype
Convergent Subtype
Age
p = 0.753
Sex
p = 1
F
M
F
M
RIN
p = 0.538
Batch
batch1
batch3
batch1
batch2
batch3
Bank
p = 0.407
ATP
NICHD
ATP
NICHD
PMI
p = 0.695
Seizures
p = 0.275
No
Yes
No
Yes
Psychiatric Medications
p = 0.447
CET
p = 0.13
ADI−R A
p = 0.589
ADI−R B(NV)
p = 0.362
ADI−R B(V)
p = 0.722
ADI−R C
p = 0.552
ADI−R D
p = 0.219

**Supplementary Figure 6**. Comparison of biological and technical covariates between the two ASD sample subtypes. Significance of categorical covariates (Cortical region, Sex, Bank, Seizures, and Psychiatric medications) was calculated using a Fisher's exact test. Significance of numerical covariates (Age, RIN, PMI, ADI-R A, ADI-R B(NV), ADI-R B(V), ADI-R C, ADI-R D, and CET) was calculated using a t-test.

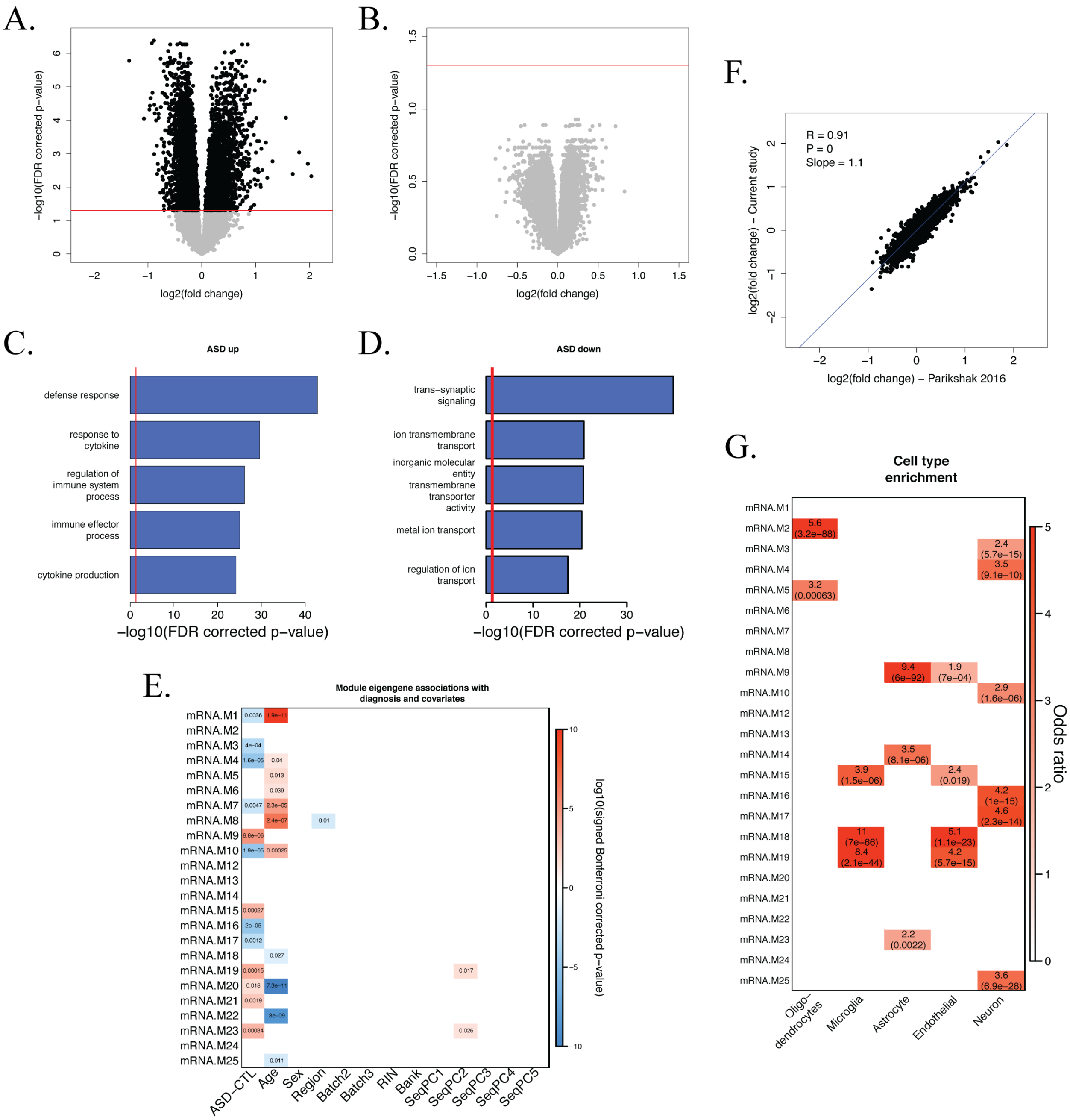
A.
−log10(FDR corrected p−value)
log2(fold change)
B.
−log10(FDR corrected p−value)
log2(fold change)
F.
R = 0.91
P = 0
Slope = 1.1
log2(fold change) − Current study
log2(fold change) − Parikshak 2016
C.
ASD up
defense response
response to cytokine
regulation of immune system process
immune effector process
cytokine production
−log10(FDR corrected p−value)
D.
ASD down
trans−synaptic signaling
ion transmembrane transport
inorganic molecular entity transmembrane transporter activity
metal ion transport
regulation of ion transport
−log10(FDR corrected p−value)
G.
Cell type enrichment
Odds ratio
Oligo−dendrocytes
Microglia
Astrocyte
Endothelial
Neuron
E.
Module eigengene associations with diagnosis and covariates
log10(signed Bonferroni corrected p−value)
ASD−CTL
Age
Sex
Region
Batch2
Batch3
RIN
Bank
SeqPC1
SeqPC2
SeqPC3
SeqPC4
SeqPC5

**Supplementary Figure 7**. mRNA expression differences in ASD.

**A**) Volcano plot showing relationship between expression fold change and significance for differential expression analysis between ASD Convergent subtype samples and controls. A significance FDR cutoff of 0.05 is shown.

**B**) Volcano plot showing relationship between expression fold change and significance for differential expression analysis between ASD Disparate subtype samples and controls. A significance FDR cutoff of 0.05 is shown.

**C**) Top gene ontology enrichments for upregulated genes in ASD.

**D**) Top gene ontology enrichments for downregulated genes in ASD.

**E**) Module-trait associations as computed by a linear mixed model with all factors on the x-axis used as covariates. Only p-values with a Bonferroni-corrected value < 0.05 are shown.

**F**) Comparison of expression fold changes between ASD Convergent subtype vs control analysis in this study and idiopathic ASD vs control analysis of Parikshak et al[9].

**G**) Cell type enrichments for all of the mRNA co-expression modules. FDR corrected p-values are shown in parentheses.

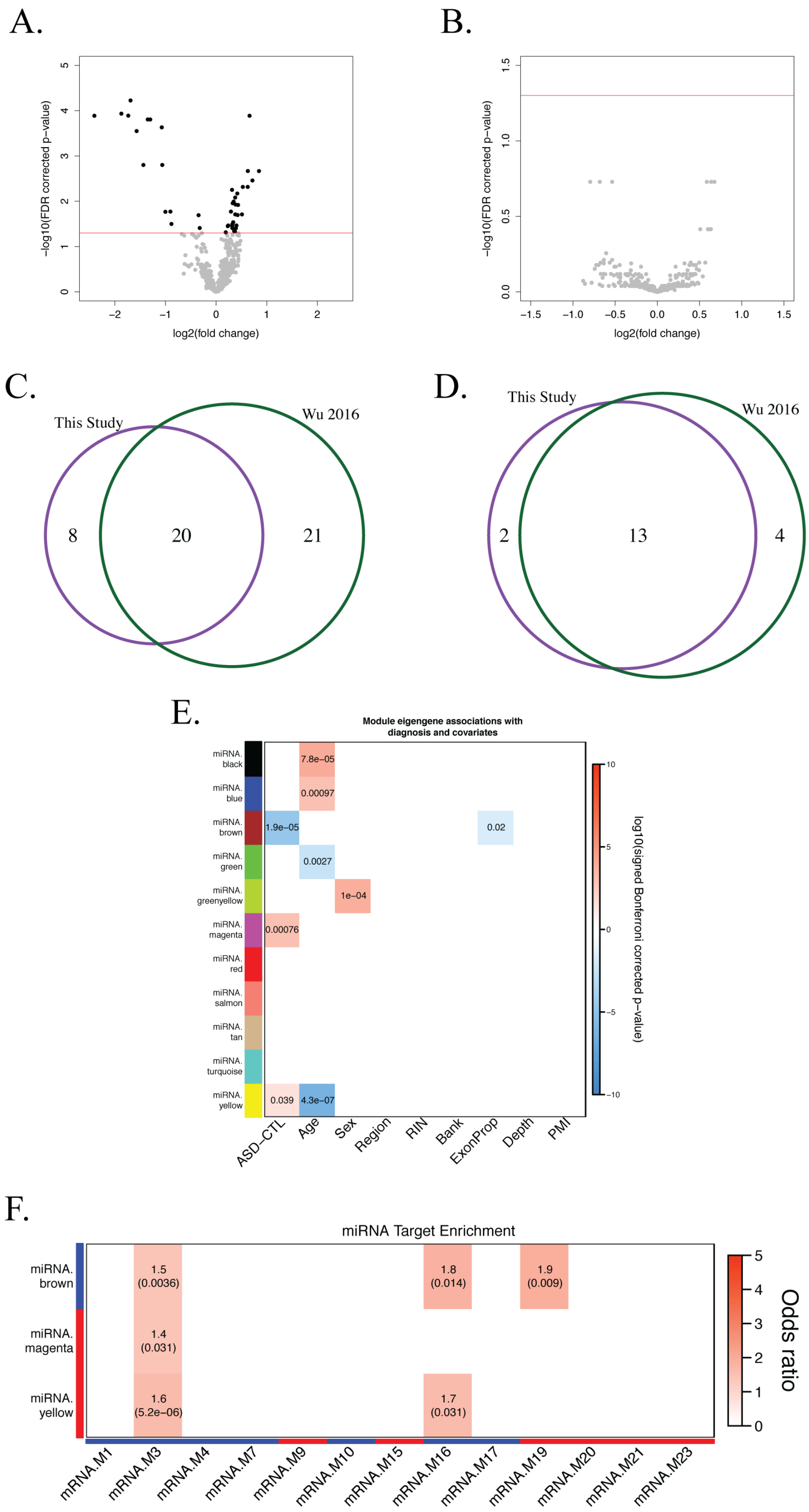
A.
−log10(FDR corrected p−value)
log2(fold change)
B.
−log10(FDR corrected p−value)
log2(fold change)
C.
This Study
Wu 2016
8
20
21
D.
This Study
Wu 2016
2
13
4
E.
Module eigengene associations with diagnosis and covariates
miRNA. black
miRNA. blue
miRNA. brown
miRNA. green
miRNA. greenyellow
miRNA. magenta
miRNA. red
miRNA. salmon
miRNA. tan
miRNA. turquoise
miRNA. yellow
7.8e−05
0.00097
1.9e−05
0.02
0.0027
1e−04
0.00076
0.039
4.3e−07
ASD−CTL
Age
Sex
Region
RIN
Bank
ExonProp
Depth
PMI
log10(signed Bonferroni corrected p−value)
F.
miRNA Target Enrichment
miRNA. brown
miRNA. magenta
miRNA. yellow
1.5 (0.0036)
1.8 (0.014)
1.9 (0.009)
1.4 (0.031)
1.6 (5.2e−06)
1.7 (0.031)
mRNA.M1
mRNA.M3
mRNA.M4
mRNA.M7
mRNA.M9
mRNA.M10
mRNA.M15
mRNA.M16
mRNA.M17
mRNA.M19
mRNA.M20
mRNA.M21
mRNA.M23
Odds ratio

**Supplementary Figure 8**. miRNA expression differences in ASD.

**A**) Volcano plot showing relationship between miRNA expression fold change and significance for differential miRNA expression analysis between ASD Convergent subtype samples and controls. A significance FDR cutoff of 0.05 is shown.

**B**) Volcano plot showing relationship between miRNA expression fold change and significance for differential miRNA expression analysis between ASD Disparate subtype samples and controls. A significance FDR cutoff of 0.05 is shown.

**C**) Overlap in miRNAs that are significantly upregulated in ASD found in this study compared to Wu et al 2016.

**D**) Overlap in miRNAs that are significantly downregulated in ASD found in this study compared to Wu et al 2016.

**E**) Module-trait associations as computed by a linear mixed model with all factors on the x-axis used as covariates. Only p-values with a Bonferroni-corrected value < 0.05 are shown.

**F**) Enrichment of differentially expressed mRNA co-expression modules with predicted targets for constituents of each differentially expressed miRNA co-expression module. Module relationships to diagnosis are marked along the x and y axes (red: increased expression in ASD; blue: decreased expression in ASD). FDR corrected p-values are shown in parentheses.

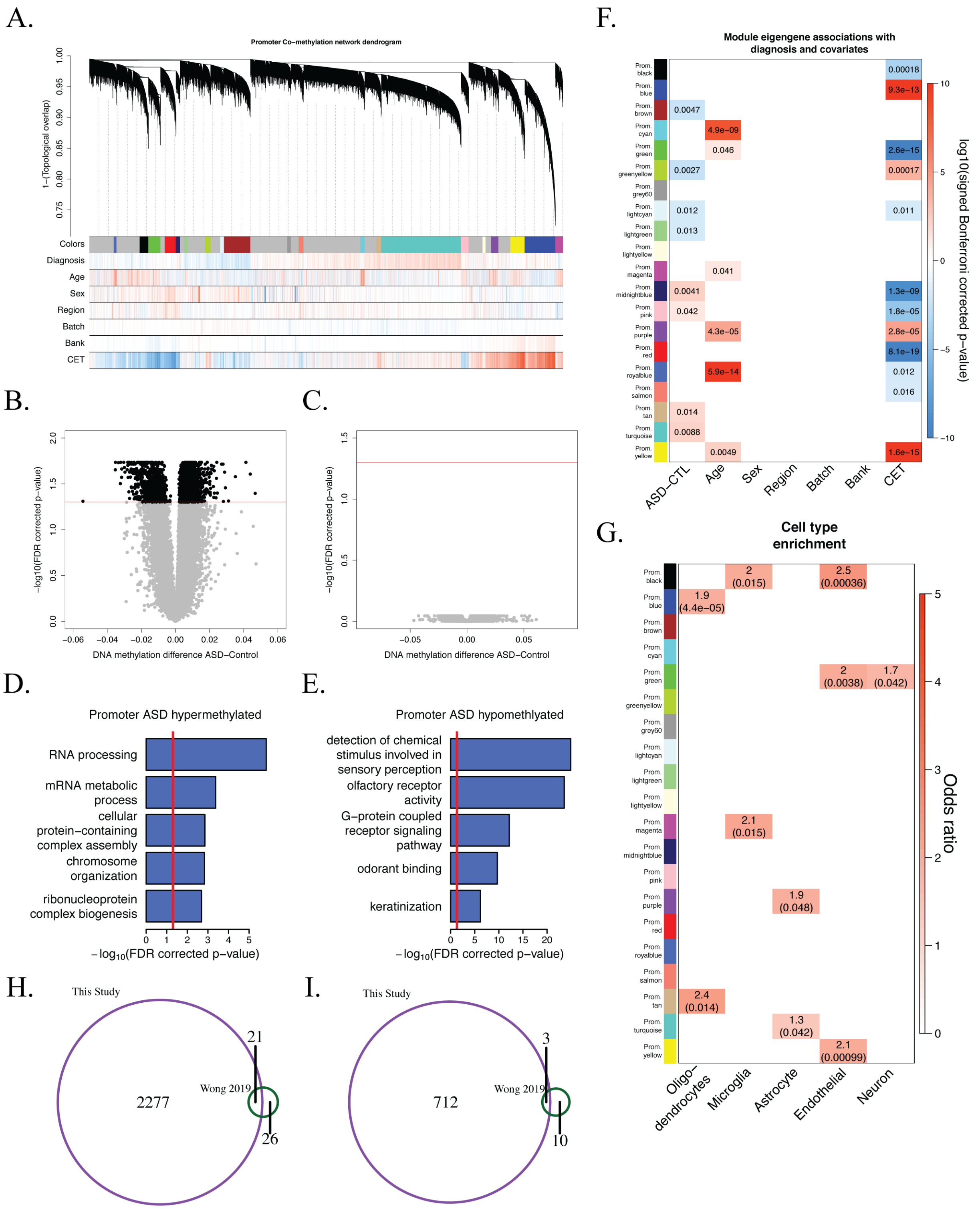
A.
Promoter Co−methylation network dendrogram
1−(Topological overlap)
Colors
Diagnosis
Age
Sex
Region
Batch
Bank
CET
B.
−log10(FDR corrected p−value)
DNA methylation difference ASD−Control
C.
−log10(FDR corrected p−value)
DNA methylation difference ASD−Control
D.
Promoter ASD hypermethylated
RNA processing
mRNA metabolic process
cellular protein−containing complex assembly
chromosome organization
ribonucleoprotein complex biogenesis
$-\log_{10}$(FDR corrected p−value)
E.
Promoter ASD hypomethlyated
detection of chemical stimulus involved in sensory perception
olfactory receptor activity
G−protein coupled receptor signaling pathway
odorant binding
keratinization
$-\log_{10}$(FDR corrected p−value)
F.
Module eigengene associations with diagnosis and covariates
log10(signed Bonferroni corrected p−value)
ASD−CTL
Age
Sex
Region
Batch
Bank
CET
G.
Cell type enrichment
Odds ratio
Oligo− dendrocytes
Microglia
Astrocyte
Endothelial
Neuron
H.
This Study
2277
Wong 2019
21
26
I.
This Study
712
Wong 2019
3
10

**Supplementary Figure 9**. Promoter DNA methylation differences in ASD.

**A**) Average linkage hierarchical clustering using the topological overlap metric for promoter co-methylation dissimilarity. Modules were identified from this dendrogram, which was constructed from a consensus of 100 bootstrapped datasets. Correlations for each gene to covariates are delineated below the dendrogram (blue, negative; red, positive). Modules are labelled with colors. The grey module represents genes that are not co-methylated and was not evaluated in further comparisons.

**B**) Volcano plot showing relationship between methylation change and significance for differential promoter methylation analysis between ASD Convergent subtype samples and controls. A significance FDR cutoff of 0.05 is shown.

**C**) Volcano plot showing relationship between methylation change and significance for differential promoter methylation analysis between ASD Disparate subtype samples and controls. A significance FDR cutoff of 0.05 is shown.

**D**) Top gene ontology enrichments for genes with hypermethylated promoters in ASD.

**E**) Top gene ontology enrichments for genes with hypomethylated promoters in ASD.

**F**) Module-trait associations as computed by a linear mixed model with all factors on the x-axis used as covariates. Only p-values with a Bonferroni-corrected value < 0.05 are shown.

**G**) Cell type enrichments for all of the promoter co-methylation modules. FDR corrected p-values are shown in parentheses.

**H**) Overlap in ASD hypermethylated gene promoters identified in this study with gene promoters containing ASD hypermethylated probes identified in Wong et al 2019.

**I**) Overlap in ASD hypomethylated gene promoters identified in this study with gene promoters containing ASD hypomethylated probes identified in Wong et al 2019.

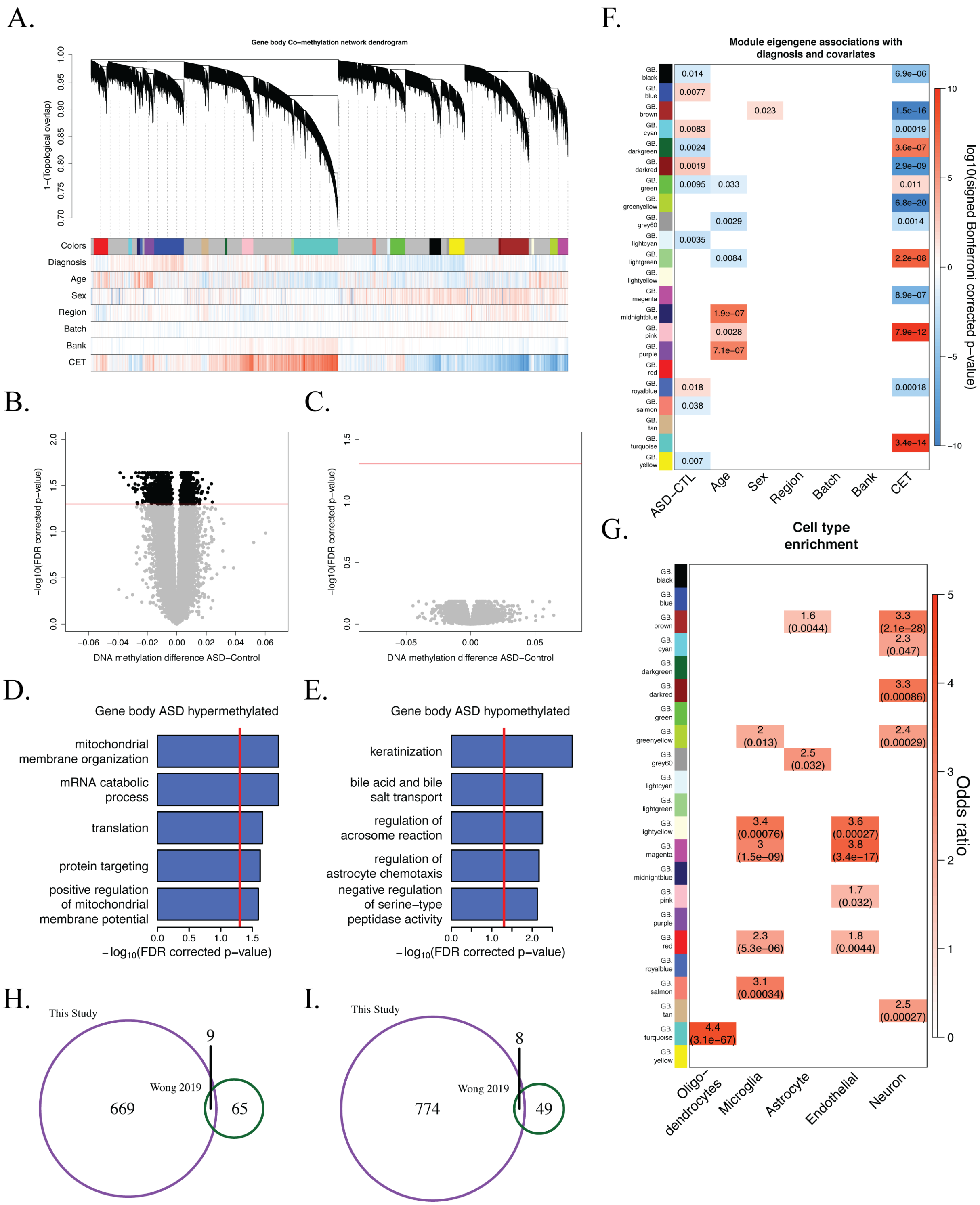
A.
Gene body Co−methylation network dendrogram
1−(Topological overlap)
Colors
Diagnosis
Age
Sex
Region
Batch
Bank
CET
B.
−log10(FDR corrected p−value)
DNA methylation difference ASD−Control
C.
−log10(FDR corrected p−value)
DNA methylation difference ASD−Control
D.
Gene body ASD hypermethylated
mitochondrial membrane organization
mRNA catabolic process
translation
protein targeting
positive regulation of mitochondrial membrane potential
−log10(FDR corrected p−value)
E.
Gene body ASD hypomethylated
keratinization
bile acid and bile salt transport
regulation of acrosome reaction
regulation of astrocyte chemotaxis
negative regulation of serine−type peptidase activity
−log10(FDR corrected p−value)
F.
Module eigengene associations with diagnosis and covariates
log10(signed Bonferroni corrected p−value)
ASD−CTL
Age
Sex
Region
Batch
Bank
CET
G.
Cell type enrichment
Odds ratio
Oligo−dendrocytes
Microglia
Astrocyte
Endothelial
Neuron
H.
This Study
Wong 2019
669
9
65
I.
This Study
Wong 2019
774
8
49

**Supplementary Figure 10**. Gene body DNA methylation differences in ASD.

**A**) Average linkage hierarchical clustering using the topological overlap metric for gene body co-methylation dissimilarity. Modules were identified from this dendrogram, which was constructed from a consensus of 100 bootstrapped datasets. Correlations for each gene to covariates are delineated below the dendrogram (blue, negative; red, positive). Modules are labelled with colors. The grey module represents genes that are not co-methylated and was not evaluated in further comparisons.

**B**) Volcano plot showing relationship between methylation change and significance for differential gene body methylation analysis between ASD Convergent subtype samples and controls. A significance FDR cutoff of 0.05 is shown.

**C**) Volcano plot showing relationship between methylation change and significance for differential gene body methylation analysis between ASD Disparate subtype samples and controls. A significance FDR cutoff of 0.05 is shown.

**D**) Top gene ontology enrichments for hypermethylated gene bodies in ASD.

**E**) Top gene ontology enrichments for hypomethylated gene bodies in ASD.

**F**) Module-trait associations as computed by a linear mixed model with all factors on the x-axis used as covariates. Only p-values with a Bonferroni-corrected value < 0.05 are shown.

**G**) Cell type enrichments for all of the gene body co-methylation modules. FDR corrected p-values are shown in parentheses.

**H**) Overlap in ASD hypermethylated gene bodies identified in this study with gene bodies containing ASD hypermethylated probes identified in Wong et al 2019.

**I**) Overlap in ASD hypomethylated gene bodies identified in this study with gene bodies containing ASD hypomethylated probes identified in Wong et al 2019.

A.

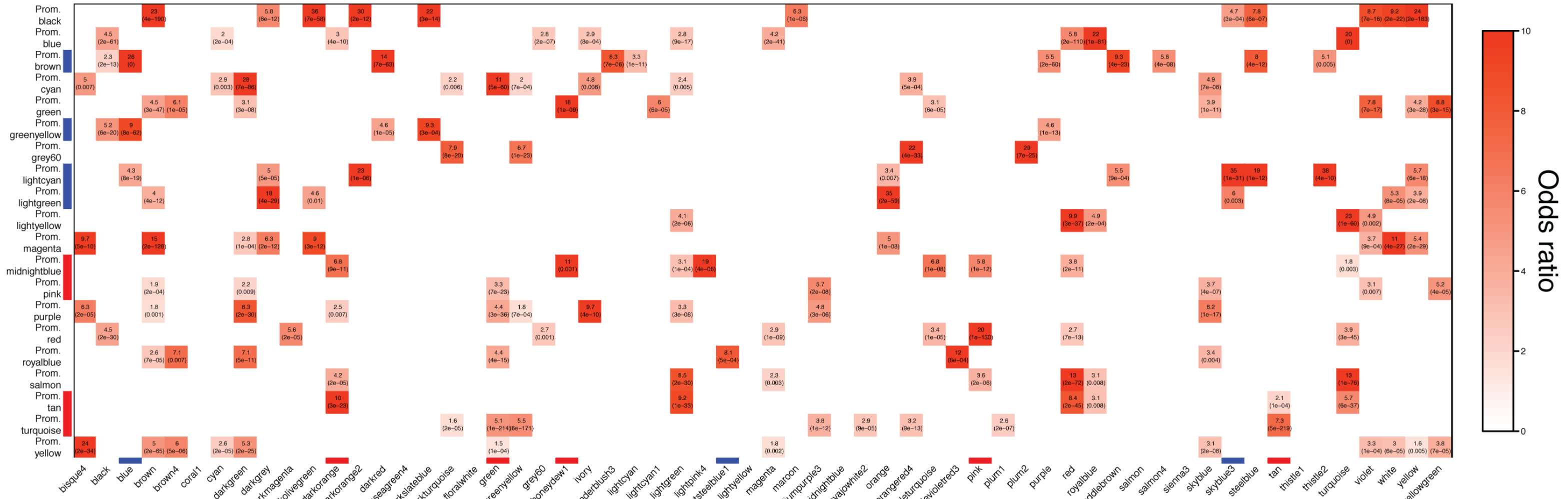

B.

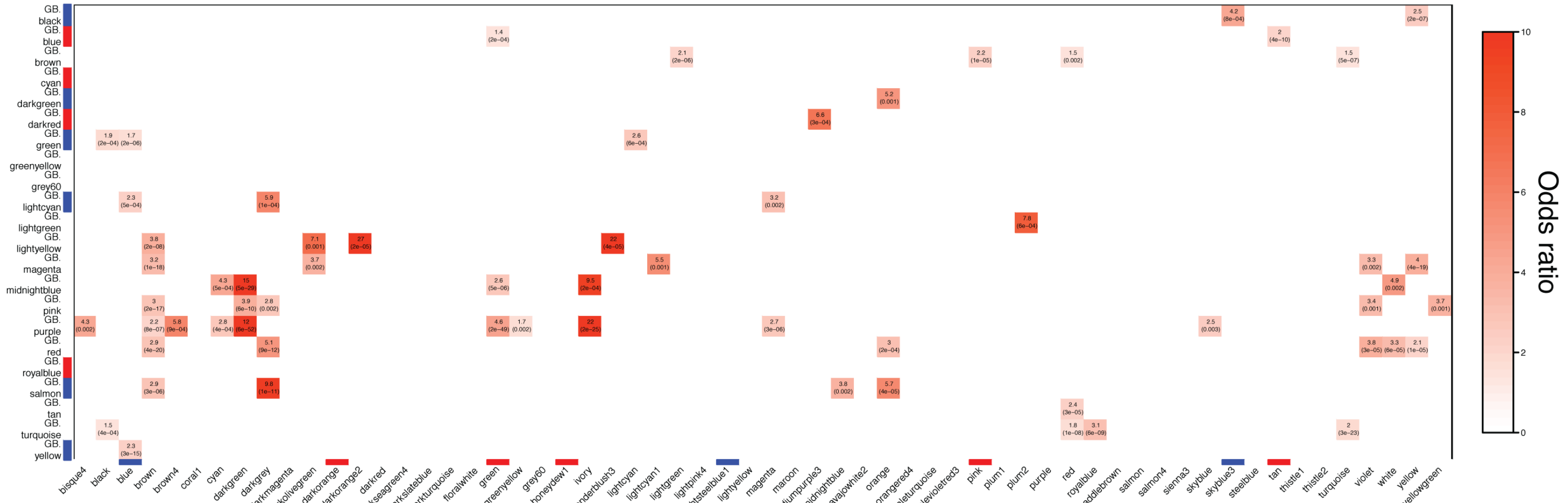

C.

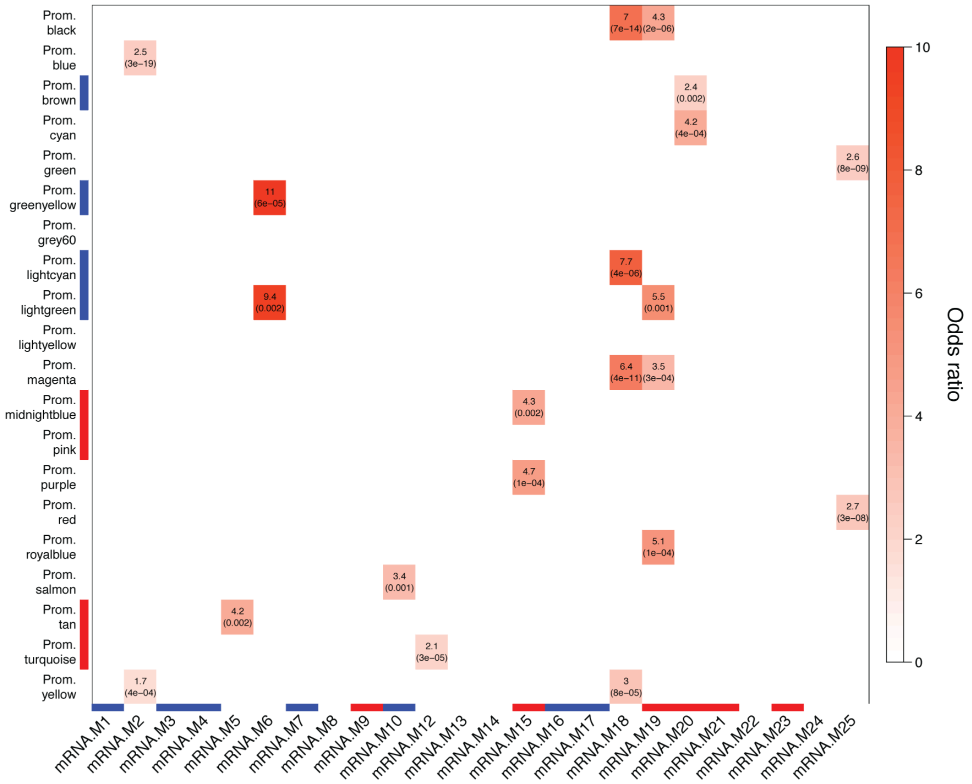

D.

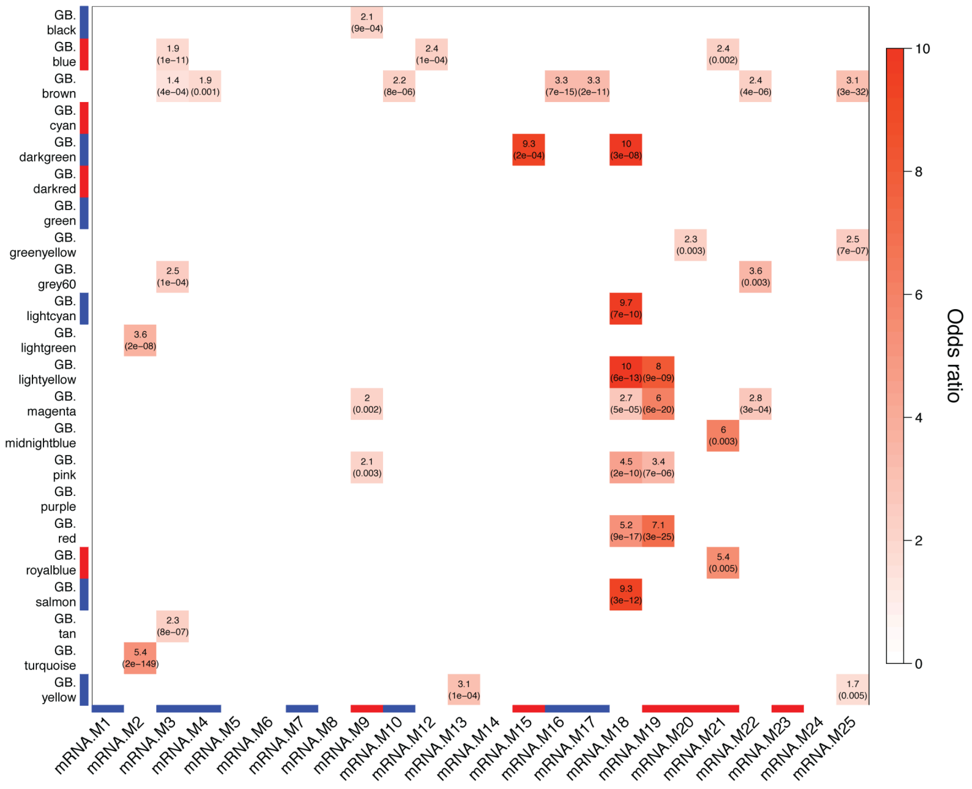

**Supplementary Figure 11**. Comparison of gene level co-methylation networks to previously published probe level co-methylation network and co-expression network.

**A**) Comparison of promoter co-methylation modules to probe level co-methylation modules identified in Wong et al 2019. Modules with a significant relationship to diagnosis are marked along the x and y axes (red: hypermethylated in ASD; blue: hypomethylated in ASD). FDR corrected p-values are shown in parentheses.

**B**) Same as **A** except for gene body co-methylation modules.

**C**) Comparison of promoter co-methylation modules to mRNA co-expression modules. Modules with a significant relationship to diagnosis are marked along the x and y axes (red: hypermethylated or increased expression in ASD; blue: hypomethylated or decreased expression in ASD). FDR corrected p-values are shown in parentheses.

**D**) Same as **C** except for gene body co-methylation modules.

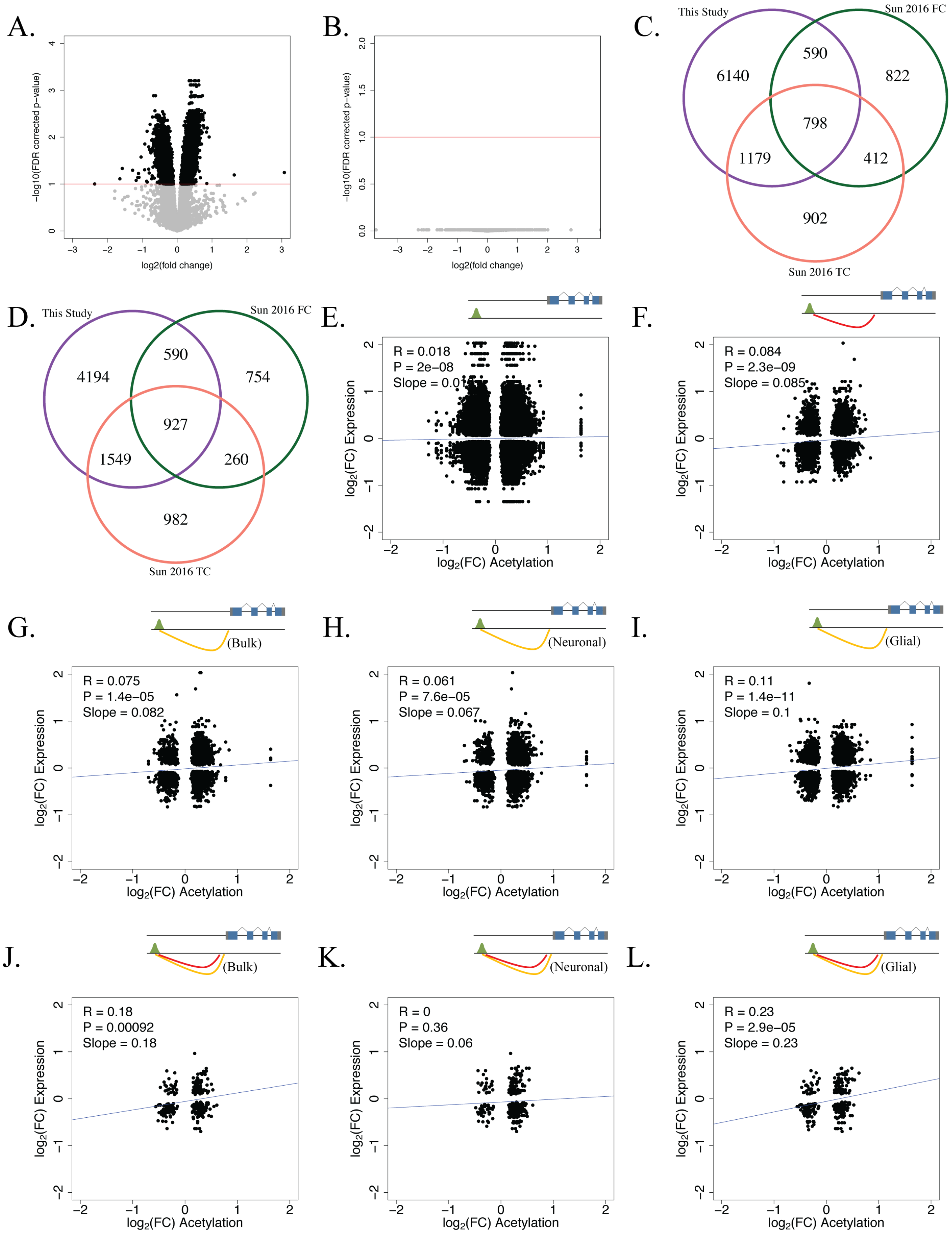

A.
−log10(FDR corrected p−value)
log2(fold change)
B.
−log10(FDR corrected p−value)
log2(fold change)
C.
This Study
Sun 2016 FC
Sun 2016 TC
6140
590
822
798
1179
412
902
D.
This Study
Sun 2016 FC
Sun 2016 TC
4194
590
754
927
1549
260
982
E.
R = 0.018
P = 2e−08
Slope = 0.01
F.
R = 0.084
P = 2.3e−09
Slope = 0.085
G.
(Bulk)
R = 0.075
P = 1.4e−05
Slope = 0.082
H.
(Neuronal)
R = 0.061
P = 7.6e−05
Slope = 0.067
I.
(Glial)
R = 0.11
P = 1.4e−11
Slope = 0.1
J.
(Bulk)
R = 0.18
P = 0.00092
Slope = 0.18
K.
(Neuronal)
R = 0
P = 0.36
Slope = 0.06
L.
(Glial)
R = 0.23
P = 2.9e−05
Slope = 0.23
log2(FC) Expression
log2(FC) Acetylation

**Supplementary Figure 12**. Histone acetylation differences in ASD.

**A**) Volcano plot showing relationship between acetylation fold change and significance for differential acetylation analysis between ASD Convergent subtype samples and controls. A significance FDR cutoff of 0.1 is shown.

**B**) Volcano plot showing relationship between acetylation fold change and significance for differential acetylation analysis between ASD Disparate subtype samples and controls. A significance FDR cutoff of 0.1 is shown.

**C**) Overlap in ASD hyperacetylated regions between this study and Sun et al 2016.

**D**) Overlap in ASD hypoacetylated regions between this study and Sun et al 2016.

**E**) Correlation between expression and acetylation changes for differentially acetylated regions within 1 MB of the TSS for differentially expressed genes. A linear model was used to correlate differential expression with differential acetylation.

**F**) Correlation between expression and acetylation changes for differentially acetylated regions linked to differentially expressed genes with eQTL evidence. A linear model was used to correlate differential expression with differential acetylation.

**G**) Correlation between expression and acetylation changes for differentially acetylated regions linked to differentially expressed genes with bulk tissue Hi-C evidence. A linear model was used to correlate differential expression with differential acetylation.

**H**) Correlation between expression and acetylation changes for differentially acetylated regions linked to differentially expressed genes with neuronal Hi-C evidence. A linear model was used to correlate differential expression with differential acetylation.

**I**) Correlation between expression and acetylation changes for differentially acetylated regions linked to differentially expressed genes with glial Hi-C evidence. A linear model was used to correlate differential expression with differential acetylation.

**J**) Correlation between expression and acetylation changes for differentially acetylated regions linked to differentially expressed genes with both eQTL and bulk tissue Hi-C evidence. A linear model was used to correlate differential expression with differential acetylation.

**K**) Correlation between expression and acetylation changes for differentially acetylated regions linked to differentially expressed genes with both eQTL and neuronal Hi-C evidence. A linear model was used to correlate differential expression with differential acetylation.

**L**) Correlation between expression and acetylation changes for differentially acetylated regions linked to differentially expressed genes with both eQTL and glial Hi-C evidence. A linear model was used to correlate differential expression with differential acetylation.

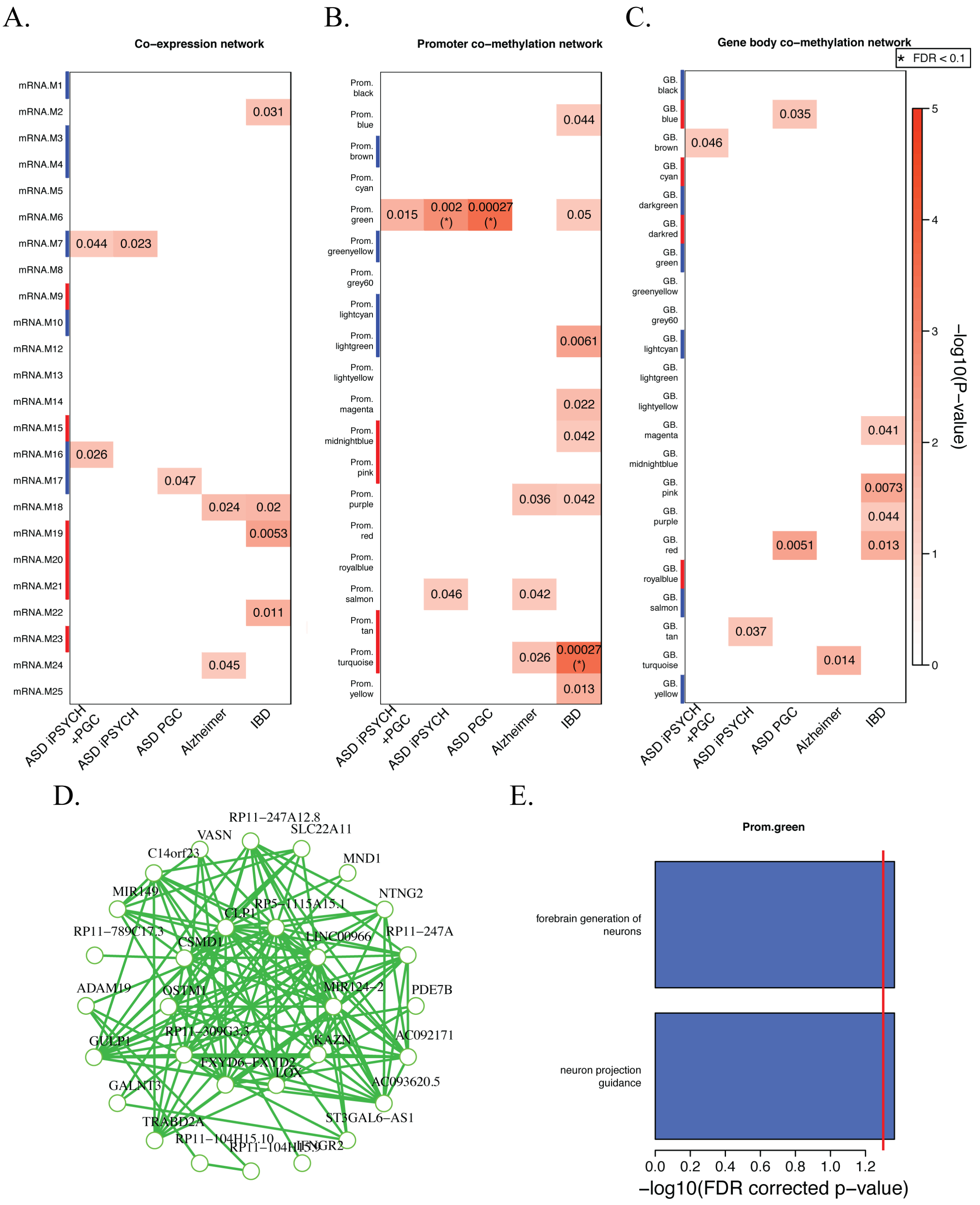
A.
Co−expression network
mRNA.M1
mRNA.M2
mRNA.M3
mRNA.M4
mRNA.M5
mRNA.M6
mRNA.M7
mRNA.M8
mRNA.M9
mRNA.M10
mRNA.M12
mRNA.M13
mRNA.M14
mRNA.M15
mRNA.M16
mRNA.M17
mRNA.M18
mRNA.M19
mRNA.M20
mRNA.M21
mRNA.M22
mRNA.M23
mRNA.M24
mRNA.M25
0.031
0.044
0.023
0.026
0.047
0.024
0.02
0.0053
0.011
0.045
ASD iPSYCH +PGC
ASD iPSYCH
ASD PGC
Alzheimer
IBD
B.
Promoter co−methylation network
Prom. black
Prom. blue
Prom. brown
Prom. cyan
Prom. green
Prom. greenyellow
Prom. grey60
Prom. lightcyan
Prom. lightgreen
Prom. lightyellow
Prom. magenta
Prom. midnightblue
Prom. pink
Prom. purple
Prom. red
Prom. royalblue
Prom. salmon
Prom. tan
Prom. turquoise
Prom. yellow
0.044
0.015
0.002 (*)
0.00027 (*)
0.05
0.0061
0.022
0.042
0.036
0.042
0.046
0.042
0.026
0.00027 (*)
0.013
ASD iPSYCH +PGC
ASD iPSYCH
ASD PGC
Alzheimer
IBD
C.
Gene body co−methylation network
★ FDR < 0.1
GB. black
GB. blue
GB. brown
GB. cyan
GB. darkgreen
GB. darkred
GB. green
GB. greenyellow
GB. grey60
GB. lightcyan
GB. lightgreen
GB. lightyellow
GB. magenta
GB. midnightblue
GB. pink
GB. purple
GB. red
GB. royalblue
GB. salmon
GB. tan
GB. turquoise
GB. yellow
0.035
0.046
0.041
0.0073
0.044
0.0051
0.013
0.037
0.014
ASD iPSYCH +PGC
ASD iPSYCH
ASD PGC
Alzheimer
IBD
−log10(P−value)
0
1
2
3
4
5
D.
RP11−247A12.8
VASN
SLC22A11
C14orf23
MND1
MIR149
NTNG2
CLPP
RP5−1115A15.1
RP11−789C17.3
CSMD1
LINC00966
RP11−247A
ADAM19
OSTM1
MIR124−2
PDE7B
GULP1
RP11−309G3.3
KAZN
AC092171
FXYD6−FXYD2
LOX
GALNT3
AC093620.5
TRABD2A
ST3GAL6−AS1
RP11−104H15.10
RP11−104H15.9
IFNGR2
E.
Prom.green
forebrain generation of neurons
neuron projection guidance
0.0
0.2
0.4
0.6
0.8
1.0
1.2
−log10(FDR corrected p−value)

**Supplementary Figure 13**. ASD genetic risk enrichments in networks.

**A**) Partitioned heritability enrichments for ASD, Alzheimer's, and IBD GWAS in mRNA co-expression modules. Modules upregulated and downregulated in ASD are marked in red and blue, respectively.

**B**) Partitioned heritability enrichments for ASD, Alzheimer's, and IBD GWAS in promoter co-methylation modules. Modules hypermethylated and hypomethylated in ASD are marked in red and blue, respectively.

**C**) Partitioned heritability enrichments for ASD, Alzheimer's, and IBD GWAS in gene body co-methylation modules. Modules hypermethylated and hypomethylated in ASD are marked in red and blue, respectively.

**D**) Top 30 hub genes and 300 connections for promoter co-methylation module Prom.green.

**E**) Top gene ontology enrichments for promoter co-methylation module Prom.green.

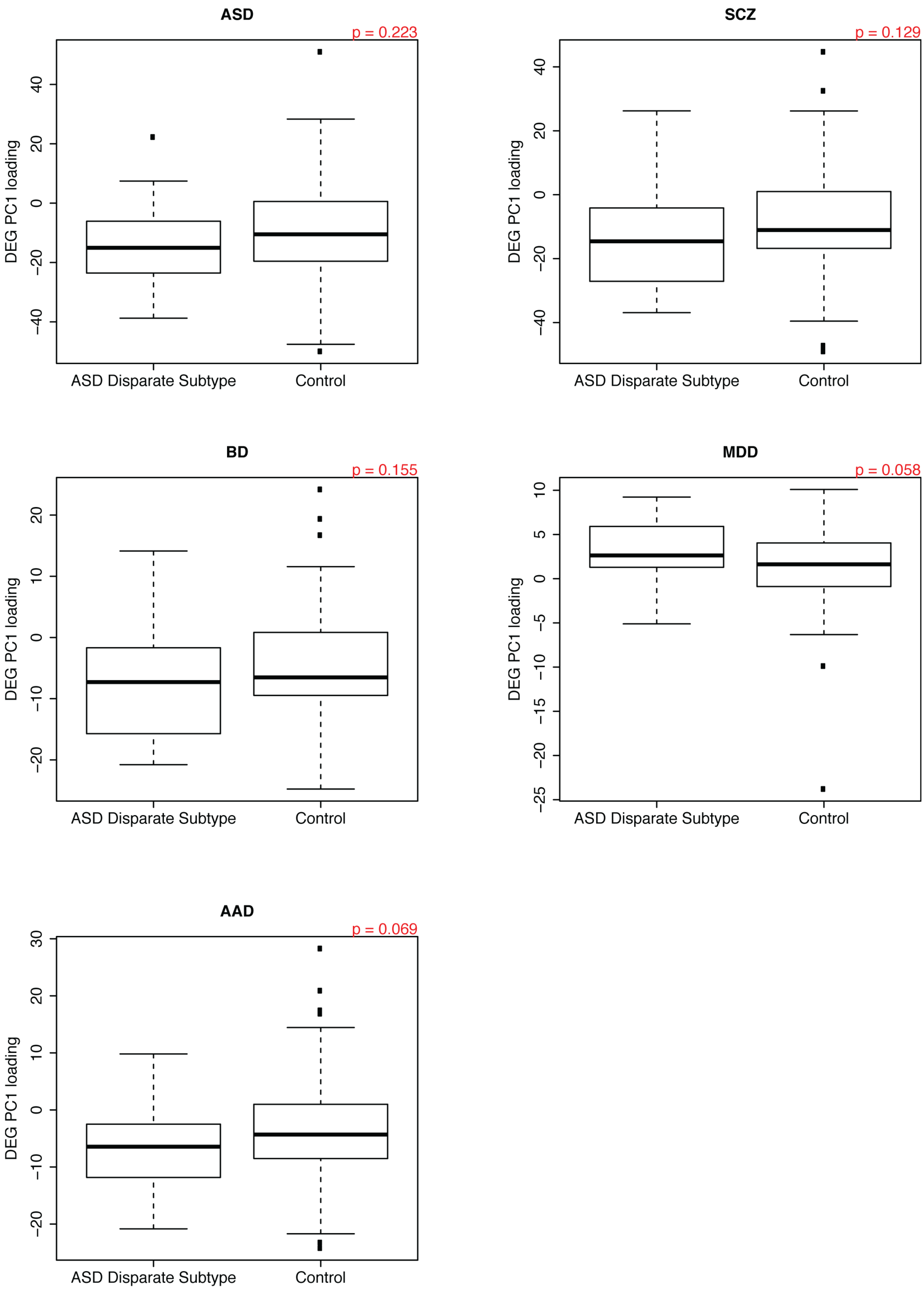
ASD
p = 0.223
DEG PC1 loading
−40
−20
0
20
40
ASD Disparate Subtype
Control
SCZ
p = 0.129
DEG PC1 loading
−40
−20
0
20
40
ASD Disparate Subtype
Control
BD
p = 0.155
DEG PC1 loading
−20
−10
0
10
20
ASD Disparate Subtype
Control
MDD
p = 0.058
DEG PC1 loading
−25
−20
−15
−10
−5
0
5
10
ASD Disparate Subtype
Control
AAD
p = 0.069
DEG PC1 loading
−20
−10
0
10
20
30
ASD Disparate Subtype
Control

**Supplementary Figure 14**. Assessment of ASD Disparate subtype samples with respect to other neuropsychiatric disorders. Loadings on the first principal component when restricting ASD Disparate subtype and control samples to differentially expressed genes from 5 neuropsychiatric disorders identified in Gandal et al 2018. P-values were calculated using a two-sided Mann-Whitney U test. ASD: autism spectrum disorder, SCZ: schizophrenia, BD: bipolar disorder, MDD: major depressive disorder, AAD: alcoholism.

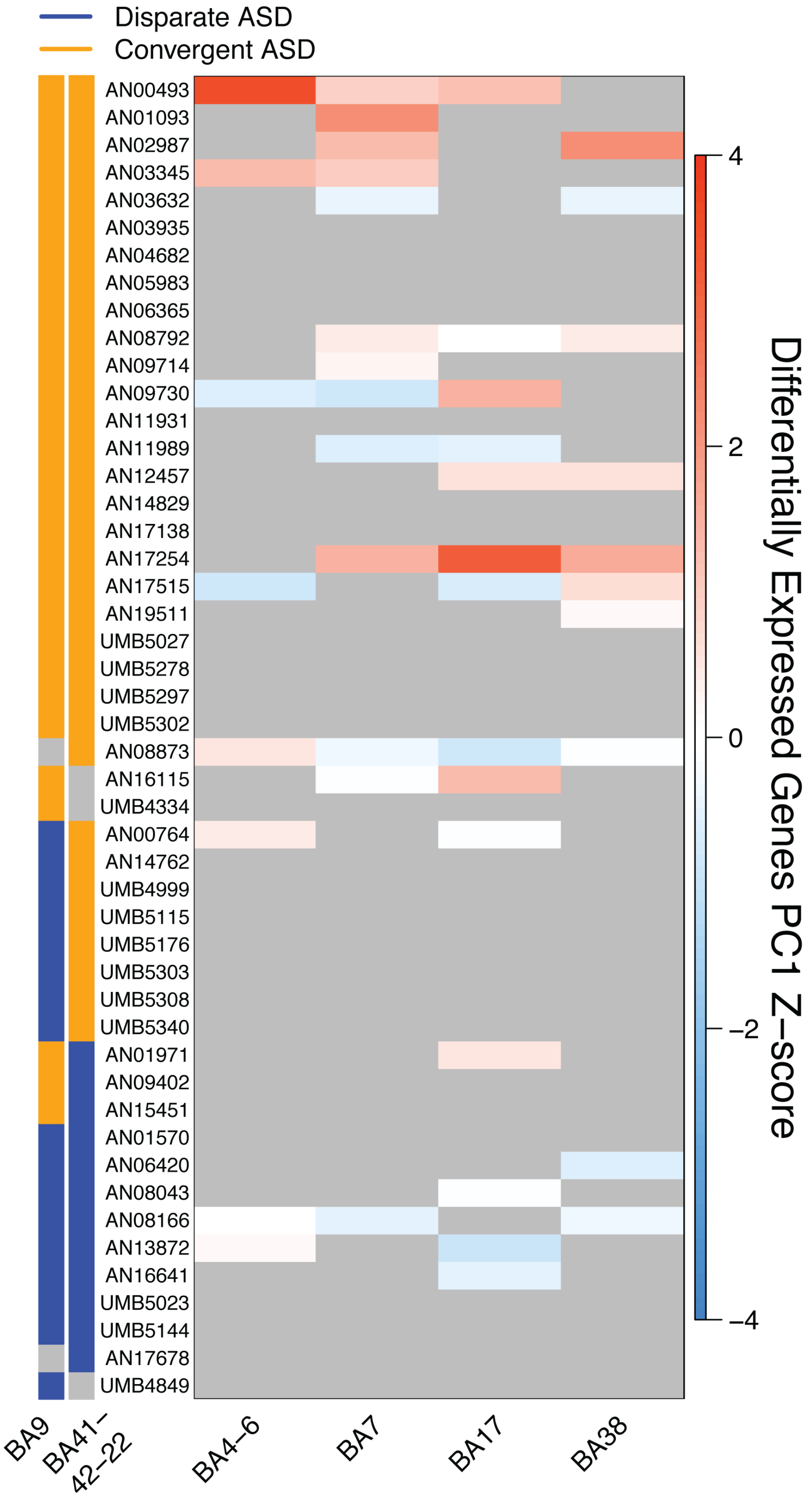

Disparate ASD
Convergent ASD
AN00493
AN01093
AN02987
AN03345
AN03632
AN03935
AN04682
AN05983
AN06365
AN08792
AN09714
AN09730
AN11931
AN11989
AN12457
AN14829
AN17138
AN17254
AN17515
AN19511
UMB5027
UMB5278
UMB5297
UMB5302
AN08873
AN16115
UMB4334
AN00764
AN14762
UMB4999
UMB5115
UMB5176
UMB5303
UMB5308
UMB5340
AN01971
AN09402
AN15451
AN01570
AN06420
AN08043
AN08166
AN13872
AN16641
UMB5023
UMB5144
AN17678
UMB4849
BA9
BA41–42–22
BA4–6
BA7
BA17
BA38
4
2
0
−2
−4
Differentially Expressed Genes PC1 Z–score

**Supplementary Figure 15**. Differential gene expression signature of ASD individuals in additional cortical regions. Loadings on the first principal component of differentially expressed genes from Parikshak et al 2016 in four additional cortical regions from Gandal et al 2018. Regions not assessed in a particular individual are colored in grey.
